# LAVENDER: latent axes discovery from multiple cytometry samples with non-parametric divergence estimation and multidimensional scaling reconstruction

**DOI:** 10.1101/673434

**Authors:** Naotoshi Nakamura, Daigo Okada, Kazuya Setoh, Takahisa Kawaguchi, Koichiro Higasa, Yasuharu Tabara, Fumihiko Matsuda, Ryo Yamada

## Abstract

Computational cytometry methods are now frequently used in flow and mass cytometric data analyses. However, systematic bias-free methodologies to assess inter-sample variability have been lacking, thereby hampering efficient data mining from a large set of samples. Here, we devised a computational method termed LAVENDER (*l*atent *a*xes disco*ve*ry from multiple cytometry samples with *n*onparametric *d*ivergence *e*stimation and multidimensional scaling *r*econstruction). It measures the Jensen-Shannon distances between samples using the *k*-nearest neighbor density estimation and reconstructs samples in a new coordinate space, called the LAVENDER space. The axes of this space can then be compared against other omics measurements to obtain biological information. Application of LAVENDER to multidimensional flow cytometry datasets of 301 Japanese individuals immunized with a seasonal influenza vaccine revealed an axis related to baseline immunological characteristics of each individual. This axis correlated with the proportion of plasma cells and the neutrophil-to-lymphocyte ratio, a clinical marker of the systemic inflammatory response. The same method was also applicable to mass cytometry data with more molecular markers. These results demonstrate that LAVENDER is a useful tool for identifying critical heterogeneity among similar, yet different, single-cell datasets.

## Introduction

Single-cell analysis is an essential approach to study heterogeneity in cell populations (1, 2). Multicolor flow cytometry is a versatile method for measuring single cells. It records expression levels of multiple surface and intracellular markers in millions of cells as they pass through the flow cell in single file while illuminated by several different lasers. Mass cytometry is a more recent technology, allowing simultaneous measurement of ∼100 markers by using heavy metal isotopes.

Currently, computational cytometry is the method of choice for analyzing high-dimensional cytometry data (3, 4). It replaced traditional analysis based on manual gates and introduced objectivity and reproducibility. However, several issues remain to be resolved.

First, the analysis is not free from the concept of gating and finding discrete cell types. For example, automatic gating methods use clustering algorithms for distinguishing different cell types, or they fit the cell distribution with predefined templates of cell types (5). Although convenient for interpretation, they fail to acknowledge quantitative variability within particular cell types that may have biological information. In fact, recent studies have even challenged the notion of discrete cell types in some cases, in favor of continuous cell states (6). In addition, predefined templates such as mixtures of Gaussians or *t*-distributions may not be adequate for highly heterogeneous cell distributions.

Second, few methods, developed so far, address the problem of comparing multiple cytometry samples systematically. Many existing approaches attempt to map each sample to the global template, which is created by pooling all samples (7–9). However, the global template, an average of samples, is both computationally demanding for large datasets and prone to neglect rare variations.

Third, even when multiple samples can be compared, there is a paucity of methods that can analyze their differences and uncover latent factors explaining sample-to-sample variability (10–13).

To solve these problems, here, we propose LAVENDER (*l*atent *a*xes disco*ve*ry from multiple cytometry samples with *n*onparametric *d*ivergence *e*stimation and multidimensional scaling *r*econstruction), a new, scalable method for comparing different cytometry samples and extracting critical axes that govern inter-sample variability. LAVENDER quantifies inter-sample differences by measuring distances between cell distributions of different samples and embedding all samples in a new coordinate space (LAVENDER space). The axes of the LAVENDER space can then be compared with other measurements for biological interpretation. It is thus an unsupervised, hypothesis-free method that can handle arbitrarily complex distributions of cells. As an application, we applied our method to peripheral blood samples from a cohort of Japanese subjects and showed that LAVENDER extracts axes of heterogeneity in the immunological states in the population. We also demonstrated that our method is applicable to mass cytometry datasets.

## Results

### LAVENDER (Latent axes discovery from multiple cytometry samples with nonparametric divergence estimation and MDS reconstruction)

LAVENDER consists of four steps (**Figure 1A**)—**Step 1**: Nonparametric density estimation of individual point clouds; **Step 2**: Distance matrix construction based on a distance metric; **Step 3**: Multidimensional scaling reconstruction of individual samples in a coordinate space; **Step 4**: Comparison of the discovered coordinates with other biological measurements.

**Figure 1.**
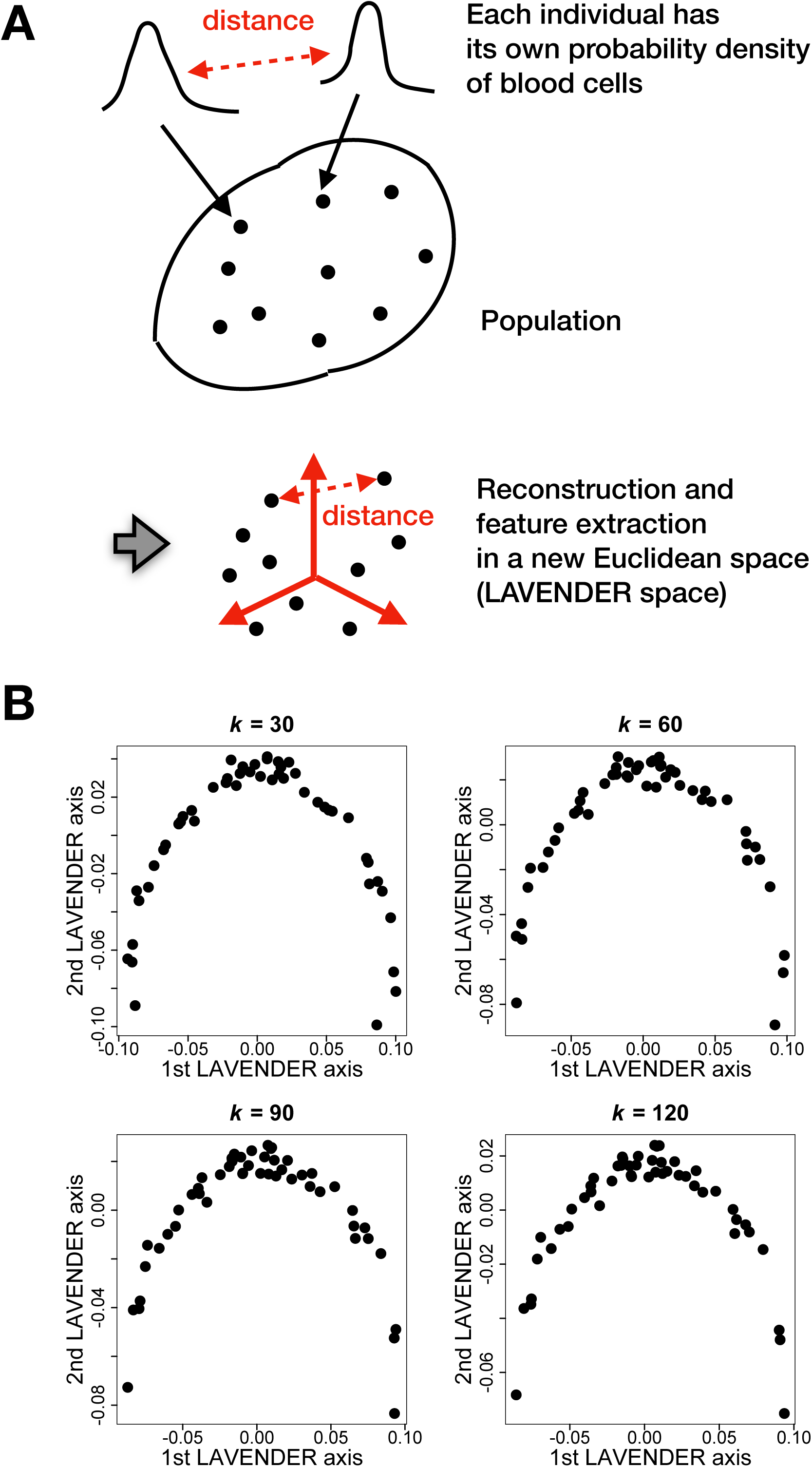
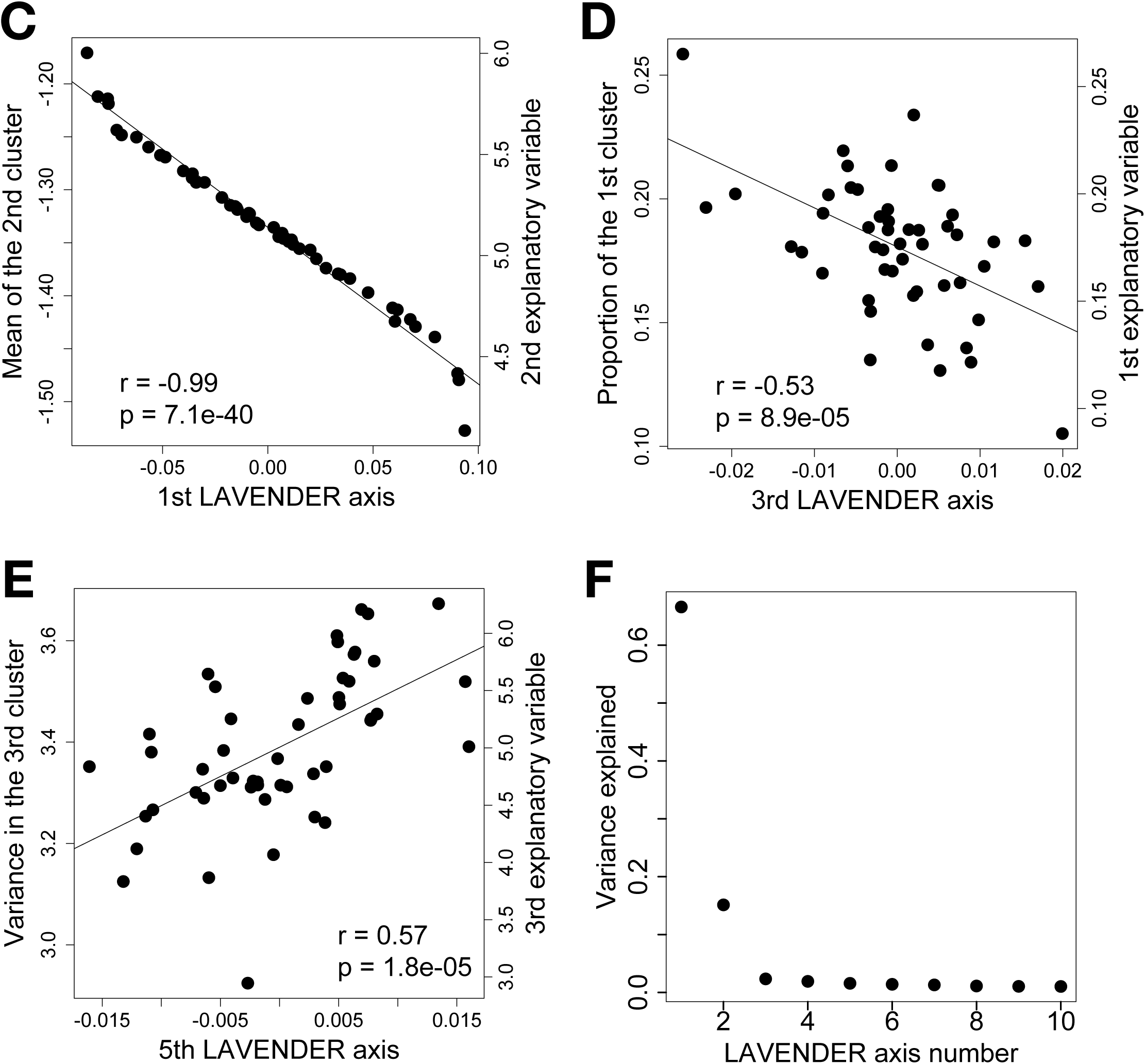
Application of LAVENDER to a synthetic flow cytometry dataset. (A) LAVENDER procedures. (B) Different values of *k* (*k* = 30, 60, 90, 120) in kNN density estimation provide similar results. (C) The first LAVENDER axis is correlated with the second explanatory variable. (D) The third LAVENDER axis is correlated with the first explanatory variable. (E) The fifth LAVENDER axis is correlated with the third explanatory variable. (F) The percentage of variance explained by each LAVENDER axis.

Each cytometry sample can be treated as a point cloud of cells in an *m*-dimensional space (cytometry space), where each point expresses the values of *m* markers measured in a single cell. We viewed each point as randomly selected from a certain probability distribution in the cytometry space. In **Step 1**, we inferred this distribution using the *k* nearest neighbor (kNN) method. Subsequently, in **Step 2**, we compared different samples by calculating the Jensen-Shannon distance between respective probability distributions. The distance reflects the difference in these distributions. In **Step 3**, based on the measured distances between all pairs of samples, we placed each sample in a new coordinate space, termed the LAVENDER space, using the algorithm of multidimensional scaling. Finally, in **Step 4**, we compared coordinates of the LAVENDER space with other biological measurements to extract biological information. Mathematical details of LAVENDER are given in the Materials and Methods section.

It is crucial not to confuse the LAVENDER space with the cytometry space. Each point in the former represents each sample containing a variety of cells, whereas that in the latter represents each cell in the sample.

### Application of LAVENDER to synthetic flow cytometry dataset

We tested LAVENDER in a synthetic dataset simulating flow cytometry. The dataset consisted of 50 samples, each containing expression levels of six markers in 10,000 cells. Every cell in a sample belonged to one of four clusters simulating different cell types. Cells in each cluster were distributed (in the cytometry space) according to a multivariate normal distribution. Three different explanatory variables, simulating biological factors, were assumed to affect the dataset. The first one increased the proportion of the first cluster and decreased that of other clusters. The second one increased the mean expression levels of markers in the second cluster. The third one increased the variance of expression levels of markers in the third cluster.

**Figure 1** shows the result of individual sample reconstruction in the LAVENDER space. Different values of *k* in kNN density estimation yielded similar results (**Figure 1B**), showing the robustness of the method. We found that the first axis in the LAVENDER space was highly correlated with the mean expression level of the first marker in the second cluster and the second explanatory variable (**Figure 1C**). The second axis was well correlated with the proportion of the first cluster and the first explanatory variable (**Figure 1D**). In addition, the fifth axis was well correlated with the variance of the first marker expression in the third cluster and the third explanatory variable (**Figure 1E**). The percentage of variance explained by each LAVENDER axis is shown in **Figure 1F**. These results demonstrate that LAVENDER can successfully extract the latent axes explaining the variability of a dataset.

### Application of LAVENDER to Nagahama flu dataset

We next applied LAVENDER to B cell samples in the Nagahama flu dataset. The dataset was obtained from peripheral blood samples of 301 Japanese participants who received a seasonal influenza vaccine (see Materials and Methods). **Figure 2** shows the result of individual sample reconstruction for Cohort A. The first three LAVENDER coordinates (*x, y, z*) of day 0, 1, 7, and 90 samples (*n* = 153, 149, 151, 148; differences are due to missing data) are shown either in 3D (**Figure 2A**) or 2D (**Figure 2B**), in black circles, red triangles, green diamonds, and blue squares, respectively. Cluster formation of same-day samples having the same color and symbol can be observed; yet, they are widely dispersed, reflecting individual variation in the immunological states. Technical replicates prepared from the same blood sample of the same subject were positioned closely in the LAVENDER space, suggesting individual variations are larger than technical variations (**Supplementary Figure 1**). LAVENDER reconstruction of Cohort B samples (day 0, 1, 7, and 90 samples, *n* = 148, 143, 147, 134; differences are due to missing data) is shown in **Figures 2C and 2D**. The percentage of variance explained by each LAVENDER axis is shown in **Figures 2E and 2F**.

**Figure 2.**
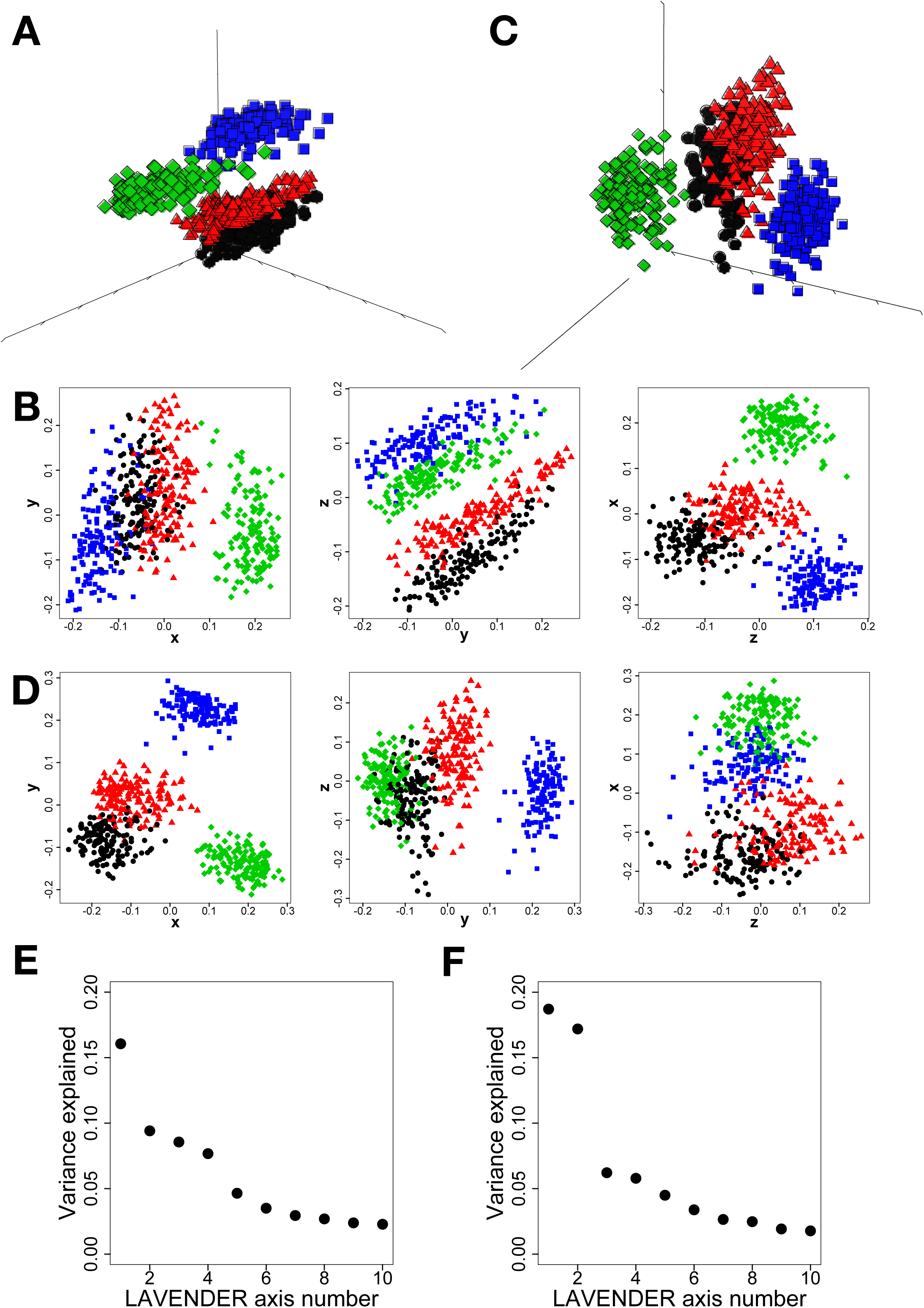
Application of LAVENDER to the Nagahama flu dataset. (A)(B) Individual sample reconstruction for Cohort A, shown in 3D (A) or 2D (B). (C)(D) Individual sample reconstruction for Cohort B, shown in 3D (C) or 2D (D). The first three LAVENDER axes are shown as *x, y*, and *z*. (E)(F) The percentage of variance explained by each LAVENDER axis. E, Cohort A; F, Cohort B.

### Individuality axis represents plasma cell proportion

The appearance of the LAVENDER space of Cohort A (**Figures 2A and 2B**) suggests that intuitively, the *x* axis represents time differences, whereas a particular direction in the *yz* plane represents individual differences. For Cohort B (**Figures 2C and 2D**), the *xy* plane represents time differences, and the *z* axis represents individual differences. To elucidate biological components constituting individual variation, we extracted the individuality axis by considering the first principal component of day 0 samples in the LAVENDER space of each cohort.

Since this axis came from our analysis of B cell samples, we hypothesized that it would be related to certain B cell subsets. Indeed, we found that, for both cohorts, the value of the individuality axis was highly correlated with the proportion of antibody-producing plasma cells in B cell samples on days 0, 1, 7 and 90, with large correlation coefficients around 0.5 and 0.7 (**Figures 3A, 4A**). This axis and the plasma cell proportion were also well correlated between different days (**Figures 3B, 3C, 4B, 4C**), suggesting they are baseline immunological characteristics inherent in each individual. However, there were no correlations between this axis and participants’ age or gender (**Figures 3D, 4D**). Comparison with clinical lab data revealed that the proportion of plasma cells was also positively correlated with the percentage of neutrophils in white blood cells (WBCs) and negatively correlated with the percentage of lymphocytes in WBCs on days 0 and 90, albeit with small correlation coefficients around 0.2 (**Figures 3E, 3F, 4E, 4F**). Taken together, the individuality axis unveiled by LAVENDER represents the plasma cell proportion, and is also weakly correlated with the neutrophil-to-lymphocyte ratio, a well-known clinical marker of systemic inflammation (14).

**Figure 3.**
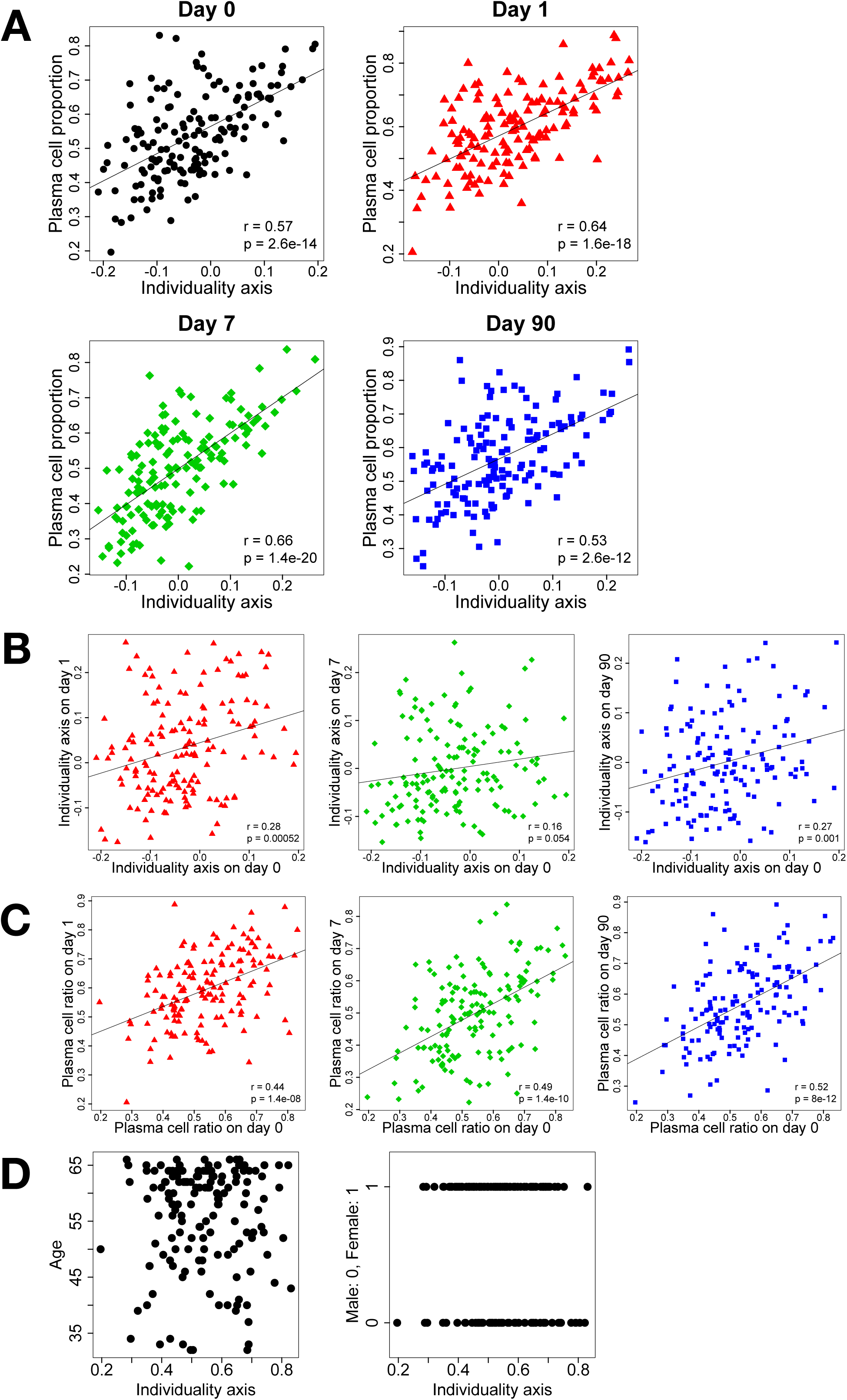
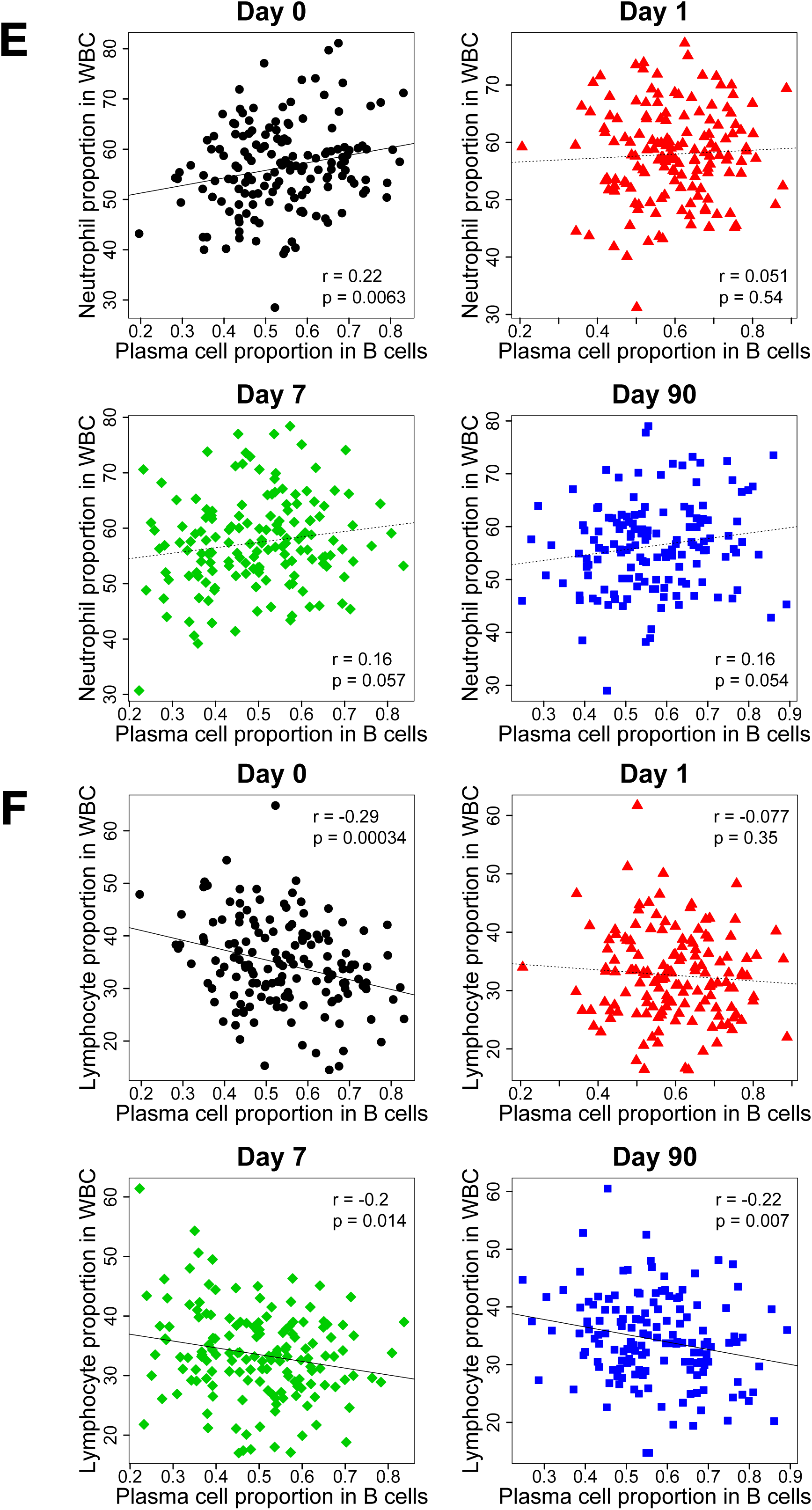
Correlation of the individuality axis with the plasma cell proportion in Cohort A. (A) The individuality axis is correlated with the plasma cell proportion. (B) The individuality axis is correlated between different days. (C) The plasma cell proportion is correlated between different days. (D) The individuality axis is not correlated with age or gender. (E) The plasma cell proportion is positively correlated with the neutrophil percentage. (F) The plasma cell proportion is negatively correlated with the lymphocyte percentage.

**Figure 4.**
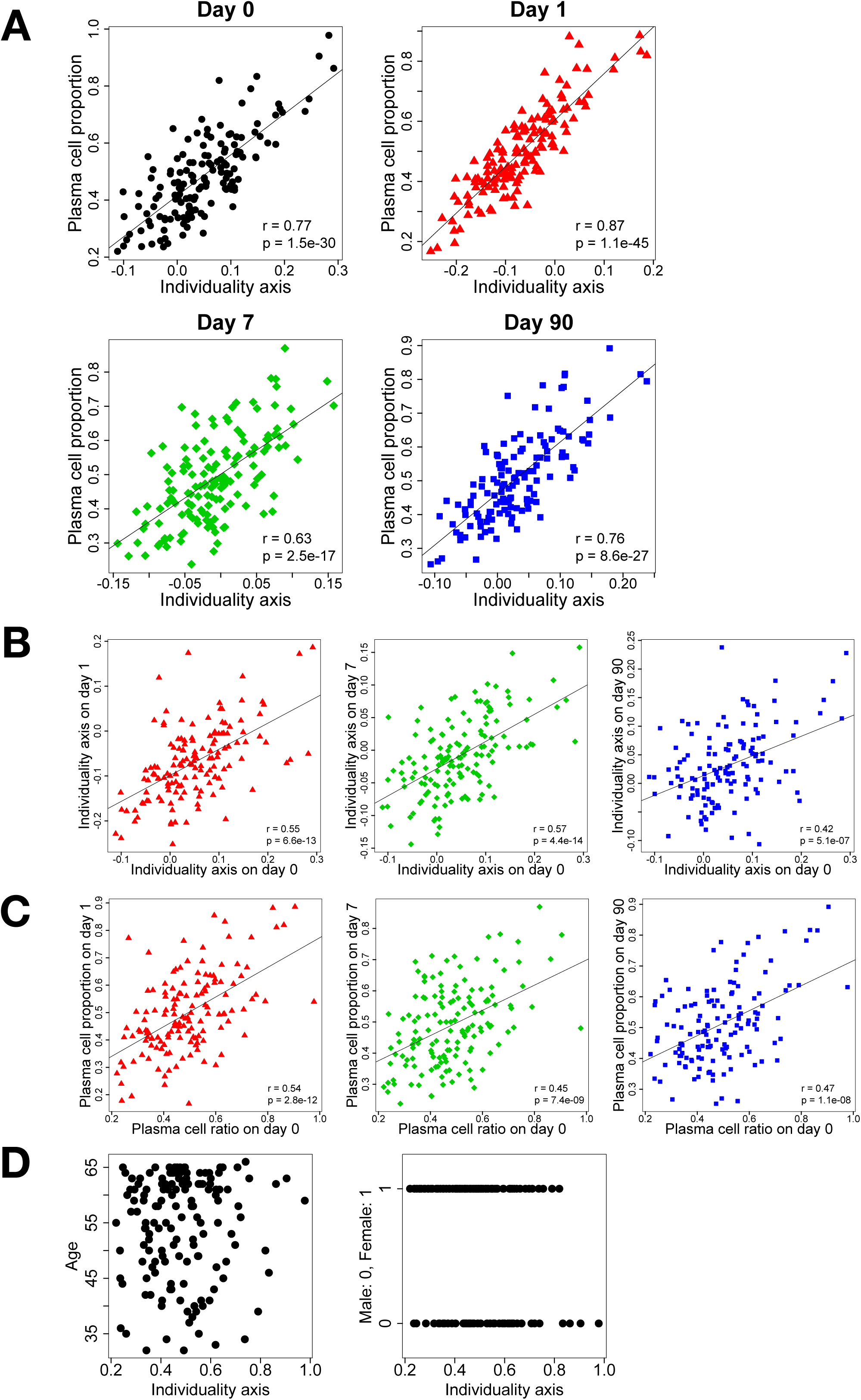
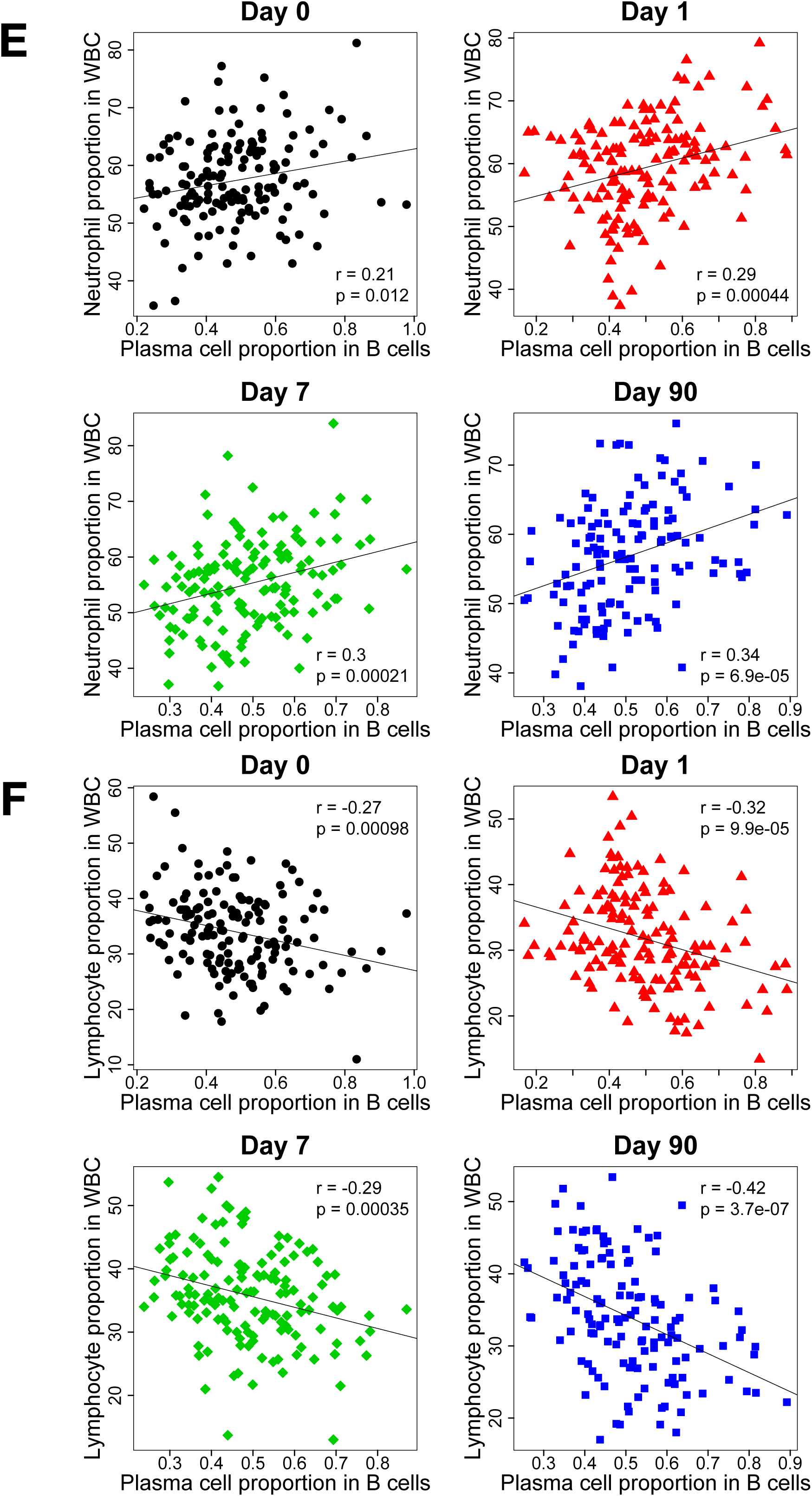
Correlation of the individuality axis with the plasma cell proportion in Cohort B. (A) The individuality axis is correlated with the plasma cell proportion. (B) The individuality axis is correlated between different days. (C) The plasma cell proportion is correlated between different days. (D) The individuality axis is not correlated with age or gender. (E) The plasma cell proportion is positively correlated with the neutrophil percentage. (F) The plasma cell proportion is negatively correlated with the lymphocyte percentage.

To further gain insight into the individuality axis, we compared against the transcriptome data of the peripheral blood. Day 0 samples were divided into two groups (Groups I and II) by means of *k*-means clustering (*k* = 2) in the LAVENDER space of each cohort (**Figures 5A, 5B**). Group I samples had larger neutrophil-to-lymphocyte ratios than Group II samples on day 0 (**Figures 5C, 5D**). Consistent with this, Gene Set Enrichment Analysis (GSEA) using Blood Transcription Modules (BTM) (15) showed that on day 0, gene sets related to neutrophils and inflammatory signaling were enriched in Group I samples, whereas those related to T cells and their activation were enriched in Group II samples (**Figure 5E**). Detailed analysis of these transcriptome data will be published elsewhere.

**Figure 5.**
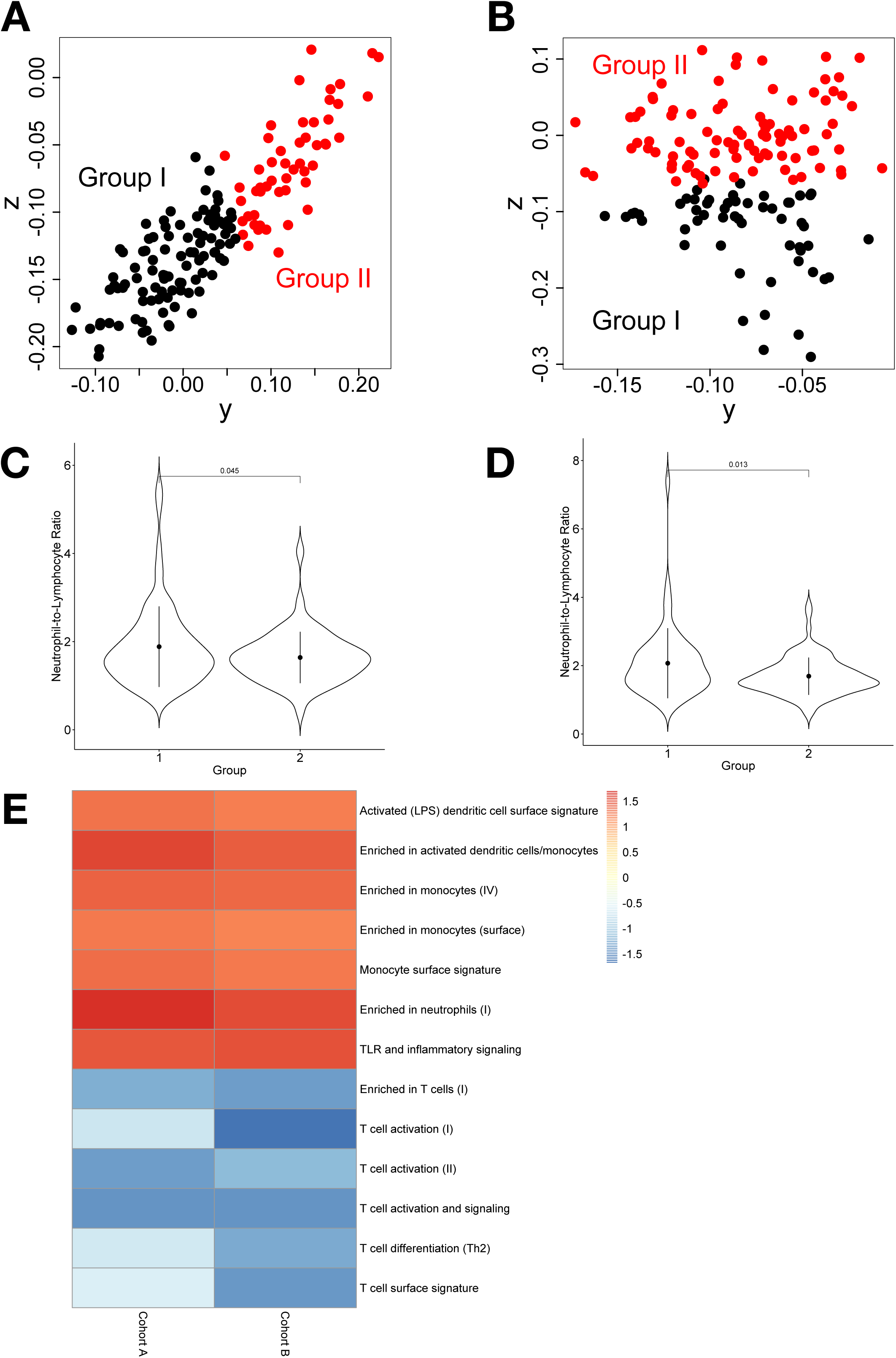
Relation of the individuality axis to the neutrophil-to-lymphocyte ratio. (A)(B) Day 0 samples are divided into Groups I and II. A, Cohort A; B, Cohort B. (C)(D) Group I samples have larger neutrophil-to-lymphocyte ratios than Group II. C, Cohort A; D, Cohort B. (E) Gene Set Enrichment Analysis using Blood Transcription Modules of Group I vs. II in both cohorts. Normalized Enrichment Scores are shown as a heatmap.

### Plasma cell proportion and HI titers

Hemagglutination inhibition assay (HI) is one of the established methods to evaluate influenza vaccine effectiveness (16, 17). Therefore, we examined the relationship between the individuality axis and HI titers. There were no significant differences between Group I and II in A/H1N1, A/H3N2, and B titers on all days (**Supplementary Figures 2A, 2B**). This might be because the plasma cell proportion is not the major determining factor of antibody titers (18), or because our titer measurement was only semi-quantitative.

We noted, however, that A/H1N1 titers on day 0 and 1 were correlated with the vaccination history of the participants (**Supplementary Figure 2C**). Specifically, those participants that were vaccinated annually had higher titers than others. Interestingly, after vaccination, the former group tended to show less increases in titers over time than others (**Supplementary Figure 2D**). As a result, A/H1N1 titers on days 7 and 90 were no longer correlated with the vaccination history, even though they were correlated with titers on days 0 and 1 (**Supplementary Figure 2E**). This trend is consistent with previous literature (19) and is possibly related to the concept of original antigenic sin (20).

### Application of LAVENDER to mass cytometry dataset

Finally, we applied LAVENDER to a public mass cytometry (CyTOF) dataset (21, 22). It measured the B cell response in peripheral blood before and after immunization with a vaccinia-based vaccine in five adult macaque monkeys over three time points (days 0, 8, 28). The result of LAVENDER construction using all 29 markers is shown in **Figure 6**. The first three LAVENDER coordinates (*x, y, z*) of day 0, 8, and 28 samples (*n* = 5, 5, 5) are shown either in 3D (**Figure 6A**) or 2D (**Figure 6B**), in black circles, green diamonds, and blue squares, respectively. The percentage of variance explained by each LAVENDER axis is shown in **Figure 6C**.

**Figure 6.**
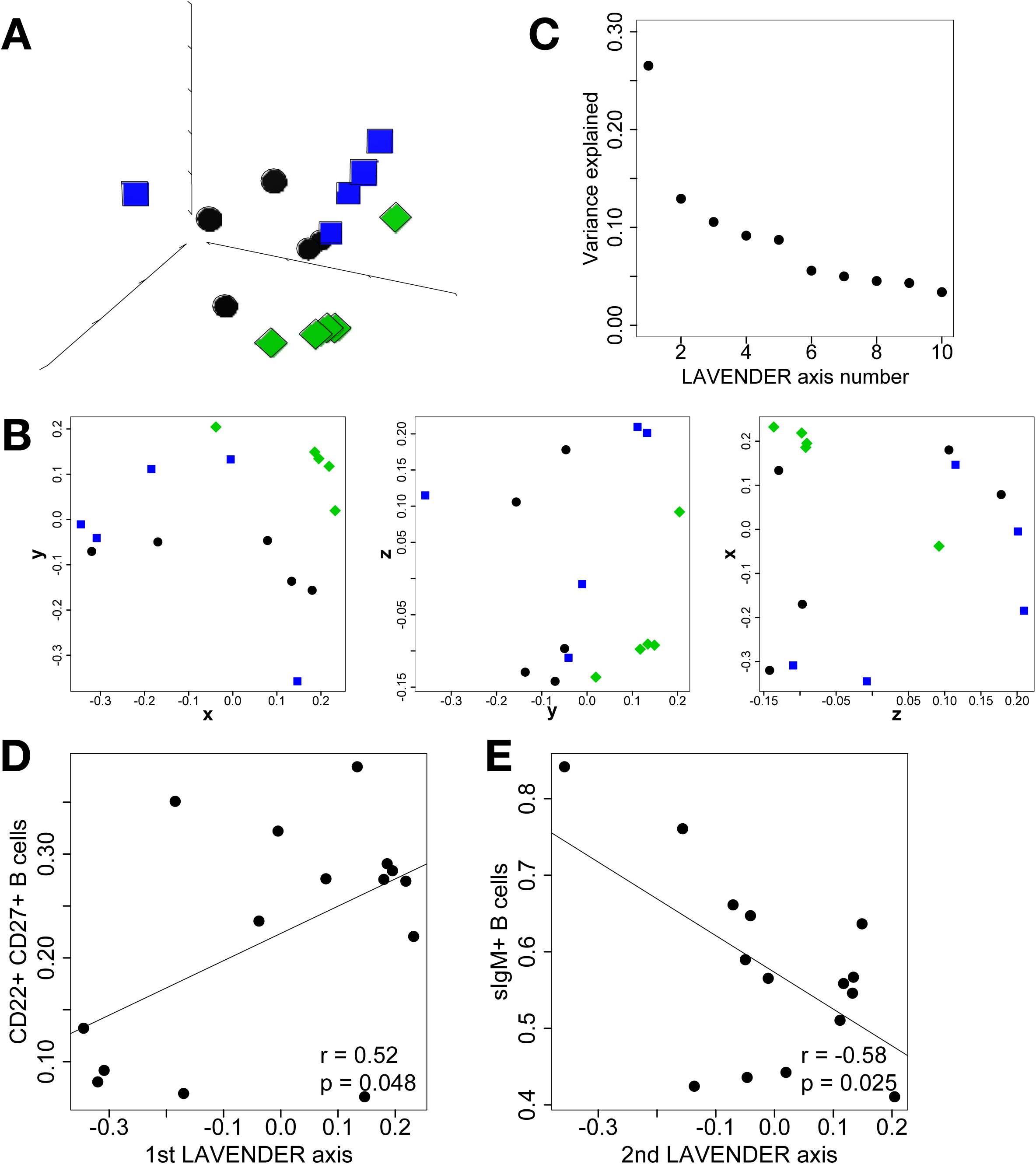
Application of LAVENDER to a public mass cytometry dataset. (A)(B) Individual sample reconstruction for Cohort A, shown in 3D (A) or 2D (B). (C) The percentage of variance explained by each LAVENDER axis. (D) The first LAVENDER axis is correlated with the proportion of CD20+ CD22+ CD27+ “memory” B cells. (E) The second axis is correlated with the proportion of sIgM+ “immature” B cells.

The appearance of the LAVENDER space suggests that the *y* axis corresponds to time differences and the *x* and *z* axes correspond to individual differences. When the LAVENDER axes were compared with B cell subsets determined by the SPADE algorithm, the *x* axis was correlated with the proportion of CD20+ CD22+ CD27+ “memory” B cells (**Figure 6D**), and the *y* axis was negatively correlated with the proportion of sIgM+ “immature” B cells (**Figure 6E**).

## Discussion

In this study, we developed a novel method of dimensionality reduction for cytometry datasets. Individual cytometry samples contain a variety of cells, each with different expression levels of surface markers. Our LAVENDER method allows comparison between different samples, summarizing them in a low-dimensional LAVENDER space, and finding latent axes of variability among the samples, all in a hypothesis-free manner. When applied to the Nagahama flu dataset, it uncovered an individuality axis and time-dependent axes. The former axis was correlated with the plasma cell proportion and the neutrophil-to-lymphocyte ratio, and our analysis suggested variability in baseline immunological states intrinsic to each individual. It was also shown that LAVENDER can be applied to mass cytometry datasets.

### Use of MDS for determination of latent axes of variability

We used a distance metric (the Jensen-Shannon distance) to quantify the difference between cell distributions of different samples. A distance metric is a distance function satisfying the triangle inequality. This facilitated the use of MDS (i.e. embedding in a Euclidean space) to visualize the result.

MDS has been utilized in a variety of biological systems as a method of clustering multiple samples (23–25). Notably, a previous study (26, 27) made use of the Gaussian kernel density estimation, the Jeffreys’ divergence (symmetrized Kullback-Leibler divergence, not a distance metric), and MDS to cluster different flow cytometry samples. They termed this method FINE (Fisher Information Nonparametric Embedding). Our LAVENDER approach, in contrast with theirs, permits the use of any distance metric not limited to approximations of the Fisher information distance. More importantly, we also demonstrated that our method can be used to discover latent axes governing the variability among samples. The discovered axes can then be compared with other multiomics data to further elucidate their biological significance.

In some choice of the distance metric, it may not be appropriate to use MDS. For example, it is known that for *p* ≠ 2, the *L*^*p*^ metric is not embeddable into the Euclidean space (28), so that the use of MDS would lead to an inexact approximation. In such cases, embedding into Riemannian manifolds (29) is a possible approach.

### Application of LAVENDER to other datasets

Although we showcased our method using flow and mass cytometry datasets, the same analysis can be extended to datasets obtained via single-cell RNA-seq (scRNA-seq) in a straightforward manner. As the number of markers (or genes) increases, the divergence estimation of cell distributions becomes more computationally demanding, mainly dependent on kNN algorithms in higher dimensional spaces. This problem can be managed using initial dimensionality reduction and/or downsampling. Initial dimensionality reduction with principal component analysis (PCA) is standard practice in the analysis of scRNA-seq data with tSNE (*t*-distributed Stochastic Neighborhood Embedding) (30, 31).

One conceptual pitfall when applying LAVENDER to scRNA-seq datasets is that in the field of single-cell biology, a sample typically means a single cell (32, 33). However, we use the same term in a different way. In our definition, a sample is a collection of cells representing a certain tissue in an individual. Currently, tissue-level scRNA-seq samples in our sense are scarce, but comparison of multiple tissue-level scRNA-seq samples, either among different individuals, tissues, or developmental stages, will become an important topic in the next several years.

## Materials and Methods

### Cytometry data

Numerical data obtained in a typical flow or mass cytometry experiment is a matrix *M*, whose rows and columns correspond to individual cells and different markers. If we have *n* cells and *m* markers, *M* is an *n* × *m* matrix and its (*i, j*) entry shows the expression level of marker *j* in cell *i*. We can display this matrix as *n* points in an *m*-dimensional Euclidean space ℝ^*m*^. The coordinates of each point show measurement results of each cell in a sample, and hereafter we identify points with cells.

We consider these measured cells to be representative of the tissue they are originally from (such as peripheral blood) and try to infer properties of the tissue from the measurement. Mathematically, we associate measured cells (points) in ℝ^*m*^ as random selections from a probability density *p*(*x*) of cells in the tissue and attempt to estimate *p*(*x*). If cells in the tissue show some meaningful tendency in terms of markers, they are expected to lie on a low-dimensional surface (manifold) embedded in ℝ^*m*^. This idea is known as the manifold hypothesis (34).

### LAVENDER (Latent axes discovery from multiple cytometry samples with nonparametric divergence estimation and MDS reconstruction)

LAVENDER consists of four steps (**Figure 1A**): (1) Nonparametric density estimation of individual point clouds; (2) Distance matrix construction based on a distance metric; (3) Multidimensional scaling reconstruction of individual samples in a coordinate space; and (4) Comparison of the discovered coordinates with other biological measurements.

**Step 1**: Each cytometry sample can be treated as a point cloud of cells in a multidimensional space (cytometry space) ℝ^*m*^, where each point *x* ∈ ℝ^*m*^ expresses *m*-channel (*m*-marker) measurement of a single cell. We view each point (cell) as a random selection from a certain probability density *p*(*x*) of cells. However, it is difficult in general to infer this probability density, because points are only sparsely positioned in the multidimensional cytometry space—a well-known phenomenon called the curse of dimensionality (35).

Nonparametric density estimation, as exemplified by the *k* nearest neighbor method (kNN), solves this problem. In kNN, a probability density *p*(*x*) around a point *x* is determined as follows. For a fixed positive integer *k*, we find a point *y* whose distance from *x* is the *k*-th smallest. We also assume that there are *n* points in total. Then, *p*(*x*) is given by

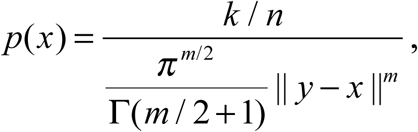

where the denominator is the *m*-dimensional volume of a sphere with radius ‖*y* − *x*‖ in ℝ^*m*^. ‖ ‖ denotes the Euclidean distance.

The benefit of using nonparametric density estimation is that we do not need to assume a particular type of distribution beforehand (as happens in parametric density estimation) and thus can flexibly express a wider variety of probability densities.

**Step 2**: After estimating probability densities for all samples in Step 1, we can quantify differences between individual samples by measuring the distance between those probability densities. From an information-theoretic point of view, the Kullback-Leibler divergence *KL* (*p* ‖ *q*), defined by

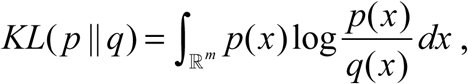

Is a natural choice for measuring the difference between probability densities *p* and *q*. For our purpose (to be described in Step 3), however, this divergence is not convenient, because it is neither symmetric

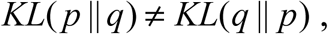

nor does it satisfy the triangle inequality

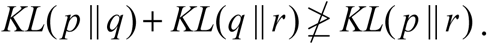

Instead, we chose the Jensen-Shannon distance, which is known to satisfy the above two conditions and is a distance metric (36):

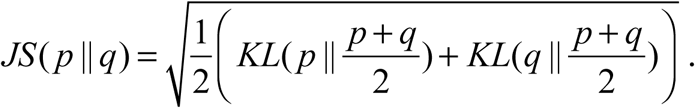

(The square of the Jensen-Shannon distance is usually called the Jensen-Shannon divergence, but to avoid confusion we do not use the latter term.) A detailed method of estimating this distance is described in the next section. It is advantageous with its close link to the Kullback-Leibler divergence, but in theory, any distance metric can be used. Intuitively, *p*(*x*) in 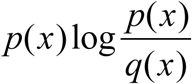 gives more weight to dense areas than sparse areas in the cytometry space, rendering it suitable for detecting biologically important differences. We also note that the Jensen-Shannon distance *JS*(*p* ‖ *q*) is bounded from above by 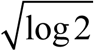, unlike the Kullback-Leibler divergence, which is not bounded from above.

**Step 3**: Based on the measured distances *d*_*ij*_ between all pairs of samples (*i, j*) (1 ≤ *i* ≤ *n*, 1 ≤ *j* ≤ *n*) in Step 2, we can reconstruct (ordinate) all samples in a new Euclidean space **ℝ** ^*K*^ (LAVENDER space), in a process called classical multidimensional scaling (MDS).

Classical MDS (also known as Torgerson MDS) is well documented (37), but we briefly explain the process below, as it is essential for understanding LAVENDER. Let 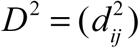 be the element-wise square of the distance matrix. We denote by *x*_*i*_ ∈ ℝ^*K*^ the position vector of each sample in the LAVENDER space and set 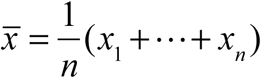. (The dimension *K* of the LAVENDER space will be determined later.) We would like to find *x*_*i*_ with ‖*x*_*i*_ − *x* _*j*_‖ as close to *d*_*ij*_ as possible.

By the law of cosines, we see that

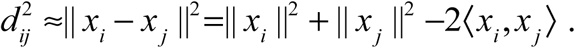

In matrix form,

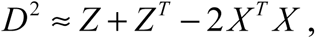

where

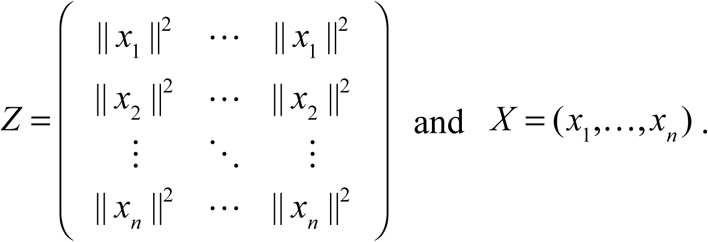

We now apply the “double centering” operation by multiplying the above by

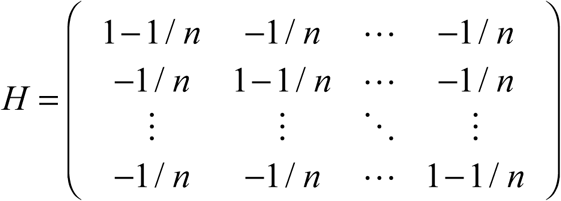

from both sides. Since *HZH* = *HZ*^*T*^ *H* = *O* and *H* = *H*^*T*^, we get

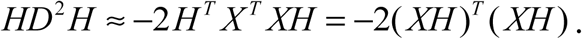

Therefore, 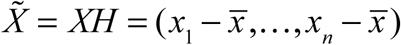 satisfies

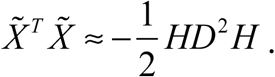

Because centering does not change the distance between samples, we can treat 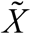 as the final coordinates of samples in the LAVENDER space. To find 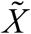, we perform eigendecomposition of the right-hand side (symmetric matrix)

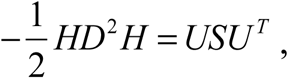

where *S* is a diagonal matrix and *U* is an orthogonal matrix. Let *S*′ be a matrix in which all negative diagonal entries of *S* (if any) are replaced by 0. Subsequently, if we set

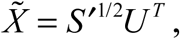

we get

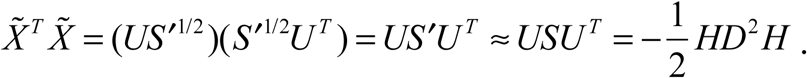

The dimension *K* of the LAVENDER space is equal to the number of positive entries in *S*. Practically, we use the space spanned by two or three eigenvectors corresponding to the two or three largest eigenvalues for visualization and later analysis.

**Step 4**: Coordinates of the LAVENDER space constructed in Step 3 can be compared with other biological measurements to extract biological information.

### Nonparametric estimation of the Jensen-Shannon distance

We now show in detail how to estimate the Jensen-Shannon distance between sample *i* and sample *j*. We first assume that sample *i* is a collection of random (i.i.d.) selections from a probability density *p*(*x*), and sample *j* is from *q*(*x*). We can then estimate the Kullback-Leibler divergence

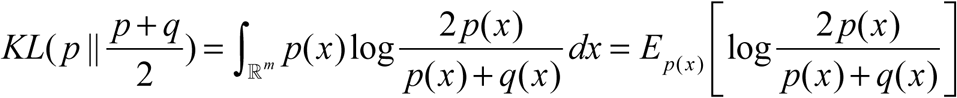

by

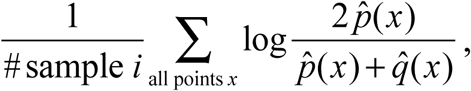

where 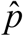 and 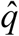 denote kNN-estimated densities of *p* and *q* at point (cell) *x*, # sample *i* denotes the number of points (cells) in sample *i*, and the summation is over all points (cells) in sample *i*. Similarly,

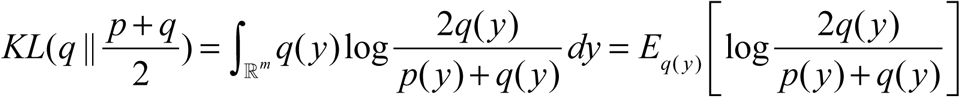

is estimated by

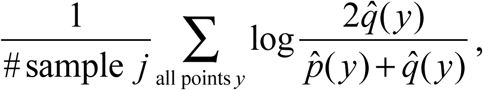

where 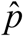 and 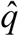 denote kNN-estimated densities of *p* and *q* at point (cell) *y*, # sample *j* denotes the number of points (cells) in sample *j*, and the summation is over all points (cells) in sample *j*. Therefore, the Jensen-Shannon distance is estimated by the nonnegative square root of

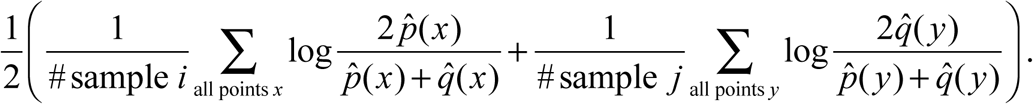

If the value of the above formula is negative, it is replaced with 0.

Note that this estimation method readily applies to other types of distance metrics, if they can be expressed as an integration of a formula containing *p*(*x*) and *q*(*x*) .

### Implementation of LAVENDER with Python and R

Flow cytometry data were preprocessed using the R package flowCore. Nonparametric density estimation using the *k* nearest neighbor method was performed either in R (using TDA and pforeach) or Python (using sklearn.neighbors and multiprocessing). Classical multidimensional scaling was carried out using the R function cmdscale. All other calculations were performed with R.

### Synthetic flow cytometry dataset

A synthetic dataset simulating flow cytometry was created to test the usefulness of LAVENDER. The dataset contained 50 samples, in which six markers were measured with six fluorescence channels. Each sample consisted of four clusters of cells, 10^4^ in total. First, the number of cells in each cluster was determined by a multinomial distribution, specified by the pre-determined proportion of each cluster. Subsequently, cells in each cluster were selected from a multivariate normal distribution in **ℝ** ^6^, specified by its mean vector and variance-covariance matrix. We further assumed five (unobserved) explanatory variables *X*_1_,…, *X*_5_, three of which influenced the dataset in the following way: *X*_1_ multiplied the proportion of the first cluster by (1+ 0.5*X*_1_), *X*_2_ multiplied the mean vector of the second cluster by (1+ 0.5*X*_2_), and *X*_3_ multiplied the variance-covariance matrix of the third cluster by (1+ 0.1*X*_3_). *X*_1_,…, *X*_5_ followed a multivariate normal distribution and were mutually independent.

### Nagahama flu dataset

The Nagahama flu dataset was obtained from a cohort of 301 Japanese volunteers challenged with a seasonal influenza vaccine. The cohort consisted of two groups (Cohorts A and B) recruited separately in Nagahama city, Shiga prefecture in western Japan. Cohort A included 100 males and 53 females, aged 32–66. Cohort B included 98 males and 50 females, aged 32–66. In winter 2011 (December 3 in Cohort A, December 17 in Cohort B), each participant had an injection of the same trivalent inactivated influenza vaccine containing three types of HA antigens from A/California/7/2009 (H1N1) pdm09, A/Victoria/210/2009 (H3N2), and B/Brisbane/60/2008. Peripheral blood samples of participants were collected before vaccination (day 0 samples), and one day, one week, three months after vaccination (day 1, 7, and 90 samples). B cell marker sets (CD19, IgM, IgD, CD21, CD27, CD138) were measured in single cells using BD FACSCanto II. Clinical lab tests were performed for the same blood samples. Hemagglutination Inhibition Assay (HI) titers for H1N1, H3N2, and B were measured according to the protocol of National Institute of Infectious Diseases, Japan. We also obtained transcriptome data for day 0 and 7 samples by extracting total RNA from the above samples and using SurePrint G3 Human GE 8×60K microarrays (Agilent #28004).

Prior to the application of LAVENDER, each cytometry sample was preprocessed by fluorescence compensation and Arcsinh transformation, followed by a B cell filter using CD19 (B cell marker) and CD138 (plasma cell marker) fluorescence levels. For each sample, a threshold value was determined by fitting the distribution of CD19 or CD138 fluorescence levels with a bimodal distribution, and all cells with lower CD19 and CD138 levels than the respective threshold were rejected. The plasma cell ratio was defined as the proportion of B cells with higher CD138 levels than the threshold.

### Public mass cytometry dataset

Raw FCS files in the mass cytometry dataset of a macaque vaccine study (22) as well as the result of the SPADE analysis were downloaded from the accompanying website of SPADEVizR (21) and analyzed according to the provided instructions. The 29 markers used were: CD20, CD69, CD3, CD38, CD197, HLADR, CD14, IgM, CD40, CD62L, CD27, CD22, Bcl-6, CD45RA, CD80, Bcl2, Ki67, CD279, IgD, B5R, CD21, CD195, CD23, CD138, IgG, CD95, CD127, TNFα, and IL10. Values in each channel were preprocessed beforehand to remove mean and unit variance.

## Acknowledgements

The authors thank Prof. James Cai, Texas A&M University, for his helpful comments on the manuscript and Dr. Maiko Narahara for her contribution in the initial phase of the study. This work was supported by JSPS KAKENHI Grant Numbers 19H05422, 17H06003, and 16KT0139 (to N.N.).

---

**Supplementary Figure 1.**
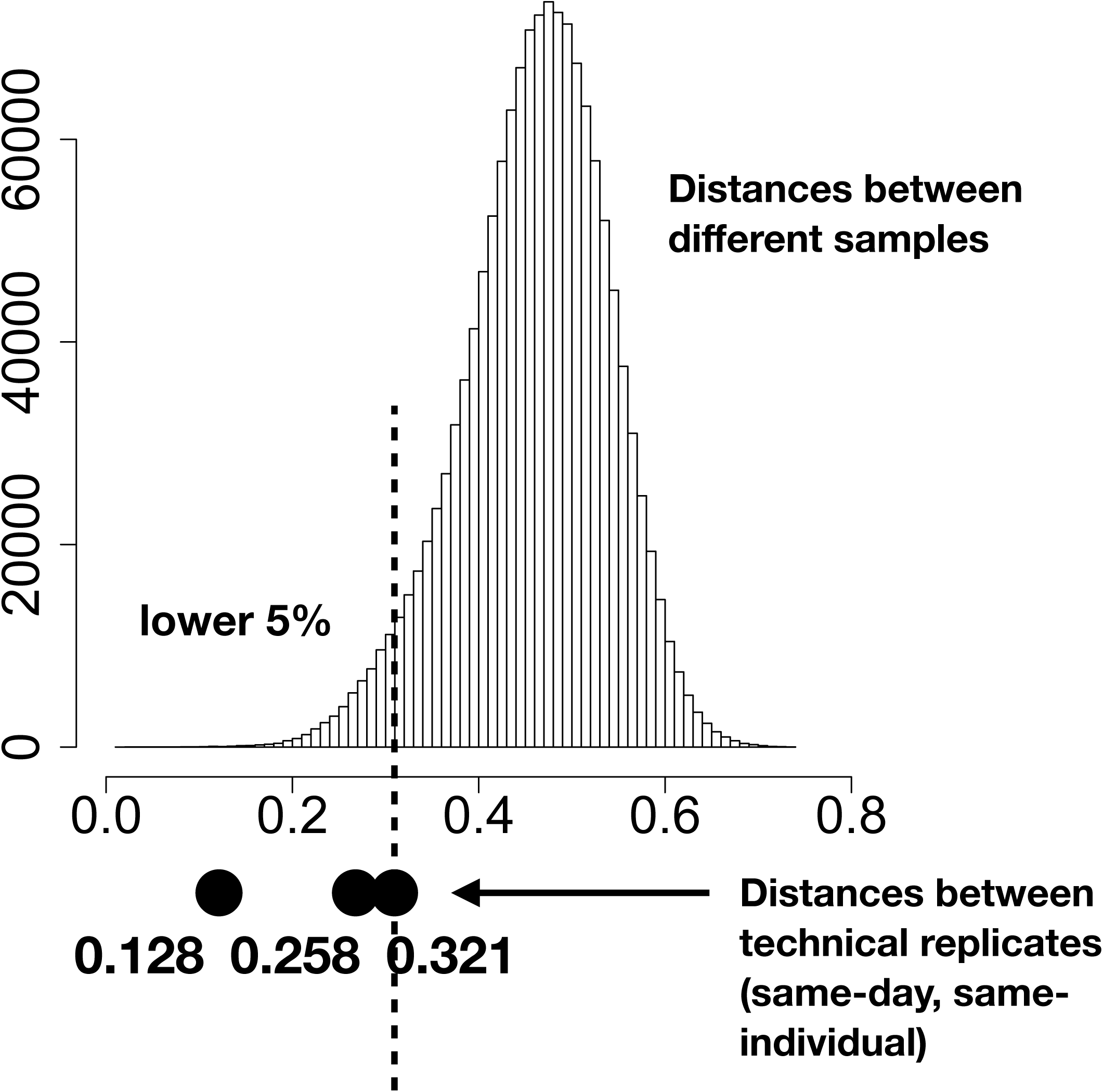
Individual variations are larger than technical variations.

**Supplementary Figure 2.**
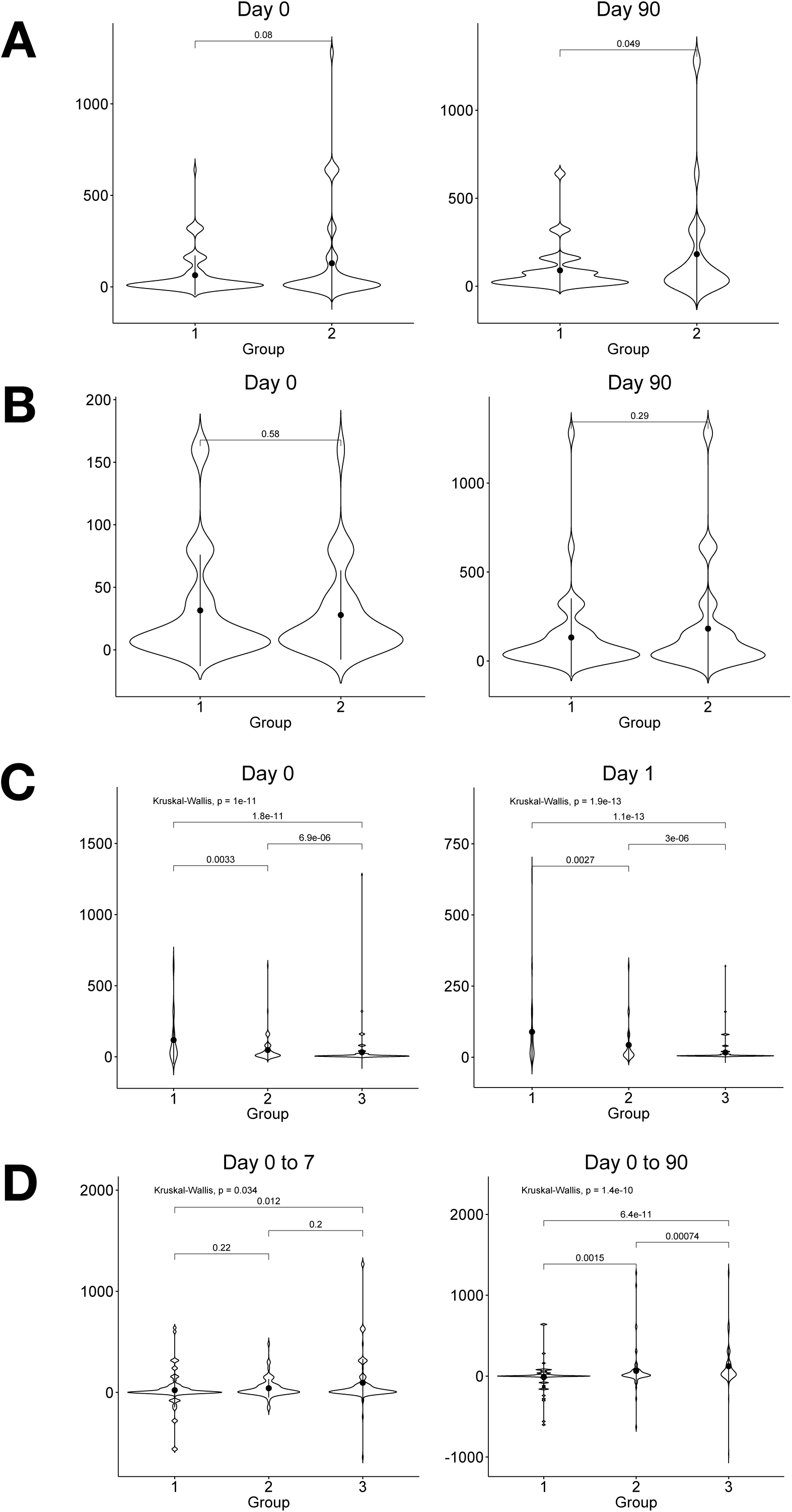
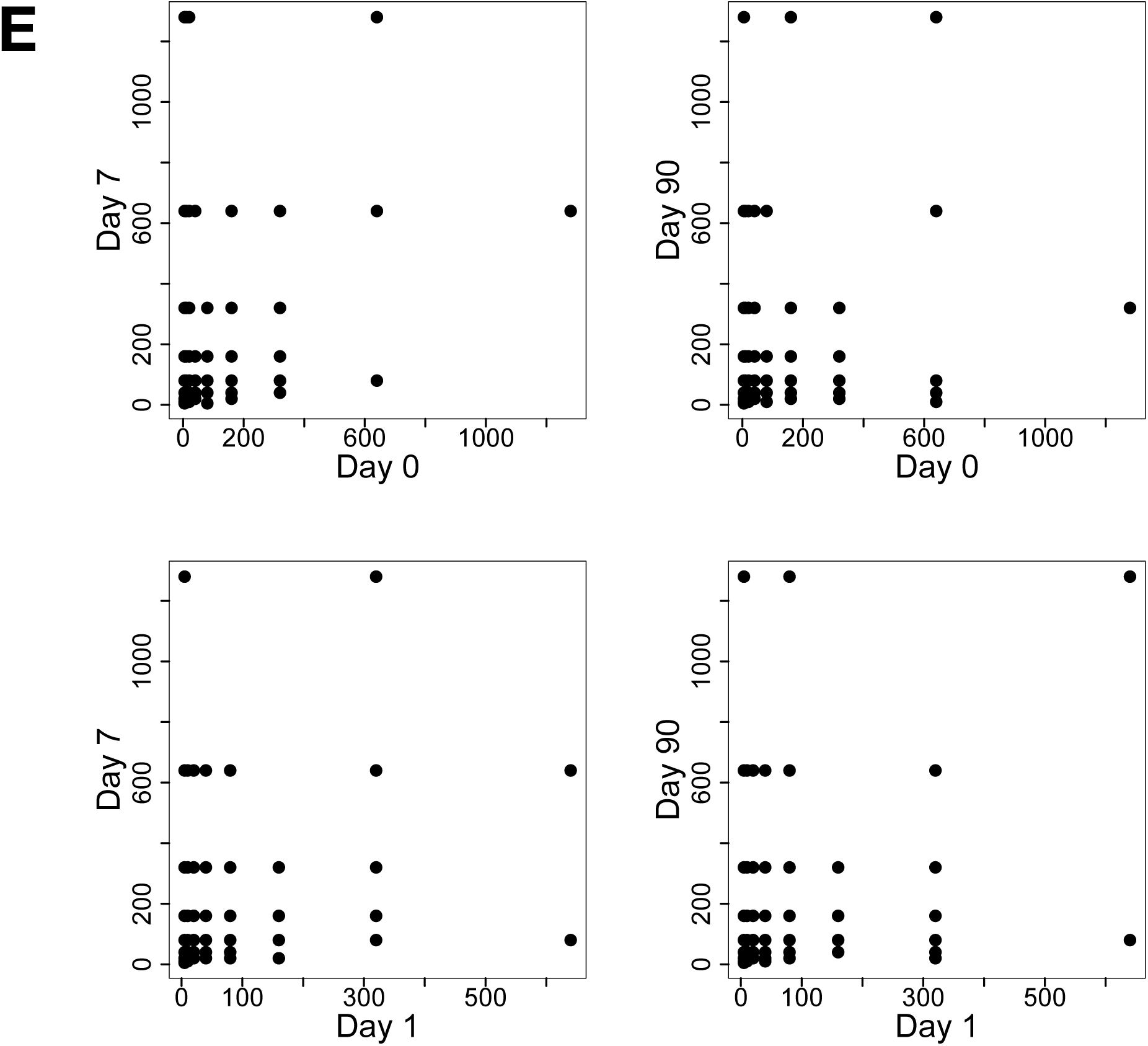
Correlation of antibody titers with vaccination history. (A)(B) A/H1N1 titers on day 0 and day 90, shown as violin plots, exhibited no differences between Group I and II. A, Cohort A; B, Cohort B. (C) A/H1N1 titers on days 0 and 1 are correlated with the vaccination history based on the participants’ questionnaire. 1, vaccinated annually; 2, vaccinated, not annually; 3, never vaccinated. (D) Change in A/H1N1 titers from day 0 to 7 or from day 0 to 90, depending on the vaccination history. (E) Correlation of A/H1N1 titers between different days.

